# DivNet: Estimating diversity in networked communities

**DOI:** 10.1101/305045

**Authors:** Amy D Willis, Bryan D Martin

## Abstract

Diversity is a marker of ecosystem health in ecology, microbiology and immunology, with implications for disease diagnosis and infection resistance. However, accurately comparing diversity across environmental gradients is challenging, especially when number of different taxonomic groups in the community is large. Furthermore, existing approaches to estimating diversity do not perform well when the taxonomic groups in the community interact via an ecological network, such as by competing within their niche, or with mutualistic relationships. To address this, we propose DivNet, a method for estimating within- and between-community diversity in ecosystems where taxa interact via an ecological network. In particular, accounting for network structure permits more accurate estimates of *alpha*- and *beta*-diversity, even in settings with a large number of taxa and a small number of samples. DivNet is fast, accurate, precise, performs well with large numbers of taxa, and is robust to both weakly and strongly networked communities. We show that the advantages of incorporating taxon interactions into diversity estimation are especially clear in analyzing microbiomes and other high-diversity, strongly networked ecosystems. Therefore, to illustrate the method, we analyze the microbiome of seafloor basalts based on a 16S amplicon sequencing dataset with 1490 taxa and 13 samples.

## 1. Introduction

Microbial communities are composed of enormous numbers of different microbes, ranging from highly abundant taxa to rare taxa that are often unobserved. Data obtained from microbiome surveys often take the form of high-dimensional count data, generally with additional covariate information regarding the experimental conditions under which the samples were observed. Detecting patterns in this data is challenging, partly because of its dimension. Analysis of *diversity* is a standard approach to summarizing and comparing high-dimensional community composition data in ecological studies, and is ubiquitous in the microbiome literature (Callahan et al. 2016). As well as providing an indicator of human and environmental health in microbiology (Oakley et al. 2008, Lozupone et al. 2012), diversity metrics are also widely used in immunology (Gibson et al. 2009, Kaplinsky & Arnaout 2016) and information theory.

Consider a community of *C* taxonomic groups (taxa), which are present in relative abundances *z* = (*z*_1_,…, *z_C_*). Depending on the ecosystem under study, *C* may be on the order of hundreds, but may also be in the tens of thousands or greater. An *α*-diversity index 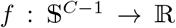 summarizes *z*, where 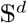 is the *d*-dimensional simplex. Similarly, *β*-diversity indices 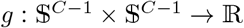 summarize information from two communities (typically, the similarity between two communities’ relative abundance vectors *z*^(1)^ and *z*^(2)^). *β*-diversity indices summarize between-community structure, while *α*-diversity indices summarize within-community structure. Specific examples of *α*- and *β*-diversity indices are given in Section 2.

Despite the prevalence of *α*- and *β*-diversity analyses in ecology, statistical methodology to estimate these functions is relatively underdeveloped. In particular, much of the existing literature focuses on estimating diversity under the assumption of observations drawn from a multinomial distribution with unknown probability vector *z* (Miller 1955, Zahl 1977, Zhang & Zhou 2010, Hsieh et al. 2016, Cao et al. 2017). Fortunately, there exist sophisticated models for community composition data that permit more a flexible co-occurrence structure than that implied by the multinomial distribution. In this paper, we use models that explicitly permit co-occurrence of taxa (commonly referred to as ecological networks) to estimate community-level diversity.

In addition to incorporating network structure, the proposed method has a number of advantages over existing methods for diversity estimation and diversity-related hypothesis testing. Most notably, while almost all existing methodology for estimating diversity either estimates the diversity of each sample (for *α*-diversity) or pairs of samples (for *β*-diversity), our method pools information across multiple samples to estimate the diversity of the ecological communities from which the samples were drawn. Therefore, rather than estimating the diversity of a sample based only on abundance information obtained from that sample, abundance information from all samples is used to improve diversity estimation. This methodology also permits a principled method for predicting diversity in ecosystems that were not sampled. Our method achieves substantial improvements in estimation performance. The method, called `DivNet`, is available as a `R` package via github.com/adw96/DivNet.

The manuscript is laid out as follows: Section 2 introduces methods for estimating *α*- and *β*-diversity. In Section 3, we introduce our model for estimating diversity, and in Section 4, we discuss estimation of the model parameters and variance estimates. The performance of the method is evaluated in Section 5, before an example of the method is discussed in Section 6. We conclude with a discussion of the method and avenues for future research in Section 7.

## 2. Literature review: Estimating *α*- and *β*-diversity

Suppose that we have samples from *i* = 1,…,*n* ecosystems. Let *C_i_* denote the set of all taxa in ecosystem *i*, and let *C_i_* = |*C_i_*| denote the number of taxa in the ith ecosystem. Let *C* = U*_i_C_i_*, and let *Q* = |*C*| denote the number of species present in one or more ecosystems. Finally, let *q* = 1,…, *Q* index the *Q* taxa. While not all taxa must be present in all ecosystems, we construct this set to ensure that the indexing is consistent. We impose the restriction that *Q* is known (see Section 7 for a discussion). Let *Z_iq_* ϵ [0, 1] denote the (unknown) relative abundance of taxon *q* in ecosystem *i*, noting that 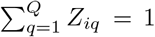. Associated with each ecosystem is a known vector of covariates 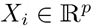.

Suppose that from the *i*th ecosystem, *M_i_* individuals are observed and classified into the *q* taxonomic groups. Let *W_iq_* denote the number of times that taxon *q* was observed in sample *i*. Therefore, to estimate summary statistics associated with the *i* ecosystems, the information available on which to base estimation is 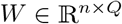 and 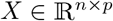.

While members of an ecological community may differ in their levels of relatedness, to constrain the scope of this paper we do not consider measures of diversity that are functions of taxonomy, such as Faith’s phylogenetic diversity (Faith 1992), branch weighted phylogenetic diversity (McCoy & Matsen 2013) or UniFrac (Lozupone & Knight 2005).

### 2.1. *α*-diversity

There are a number of different *α*-diversity indices that are widely used in the literature. This is because different indices reflect different features of ecosystems. Two of the most common indices are the Shannon entropy (also called the Shannon index), and the Simpson index. The Shannon index places more emphasis on rare species than the Simpson index (because – *x* log *x* μ *x*^2^ for *x* close to zero; see Eqs. (1) and (5)). Therefore, in ecosystems where rare species are significant drivers of ecosystem health, the Shannon index may be preferred over the Simpson index (for example, see Oakley et al. (2008)). While the diversity estimation framework that we will introduce is applicable to any *α*-diversity index that is a function of taxon abundance, we will focus on the Shannon and Simpson indices to illustrate our method.

#### 2.1.1 Shannon entropy

One of the most common *α*-diversity indices is the Shannon entropy (Shannon 1948). The Shannon index of ecosystem *i* is defined as

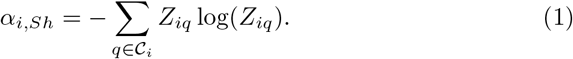

This index captures information about both the species richness (number of species) and the relative abundances of the species. Specifically, as the number of species in the population increases, so does the Shannon index. As the relative abundances diverge from a uniform distribution (*Z_iq_* = 1/*C_i_* for all *q* ϵ *C_i_*) and become more unequal, the Shannon index decreases: for fixed |*C_i_*|, the entropy is maximized when the abundance of all taxa is equal.

Under the model **W***_i_* ∼ *Multinomial*(*M_i_*, **Z***_i_*), the maximum likelihood estimate (MLE) of *α_i,Shannon_* is

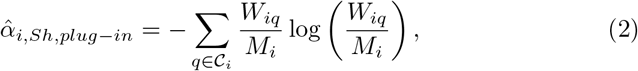

with the convention that if *W_iq_* = 0, then 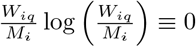, since lim*_x⟶0_ x* log *x* = 0. This estimate is almost ubiquitous in the ecological literature (Weiss et al. 2017, Willis 2017). The multinomial MLE is often referred to as the *plug-in* estimate (Vu et al. 2007). The multinomial MLE is negatively biased by 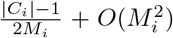 (Basharin 1959), for which various corrections have been proposed, including adding 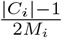 (the *Miller-Maddow* MLE correction, Miller (1955)), and jackknifing (Zahl 1977).

Noting that unobserved (latent) taxa are often a substantial source of error in estimating the Shannon index, Chao & Shen (2003) proposed using the Good-Turing estimate of species richness and adjusting for the missing taxa, obtaining the estimate

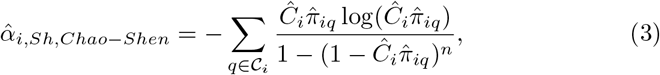

where 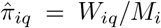 and 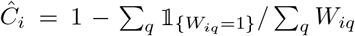. Vu et al. (2007) show that this estimator is consistent and converges with the optimal rate *Op* (1/log(*M_i_*)).

More recently, Chao et al. (2013) proposed to correct bias due to latent taxa by subsampling taxa and extrapolating from the sequentially smaller subsamples. The method is implemented in the R package `iNEXT` (Hsieh et al. 2016), against which we compare our method. We note that the subsampling procedure of `iNEXT` involves subsampling the taxa independently, which reflects the assumptions of the multinomial model.

An alternative approach to adjusting for latent taxa originates in the compositional data analysis literature. To estimate the compositions *Z_iq_*, Martín-Fernández et al. (2003) propose replacing observed values of *W_ij_* that are exactly zero with 0.5, and so Cao et al. (2017) consider the resulting *zero-replace α*-diversity estimator

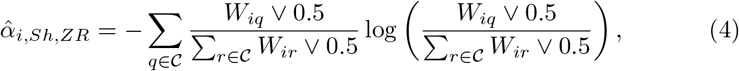

and also extend this idea to imputing zero elements of *W* via a low-rank matrix projection using a regularization approach based on a Poisson-Multinomial model. No publicly available software implements the low-rank matrix method.

#### 2.1.2 Simpson index

Simpson (1949) defined the index now known as the *Simpson index:*

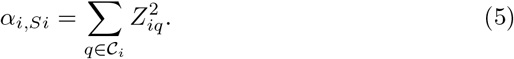

Similar to the Shannon index, the most common estimate of the Simpson index
is the plug-in estimate:

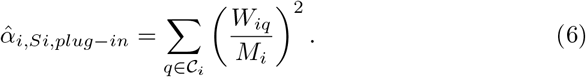

In comparison with Shannon entropy estimation, research concerning optimality of estimates of the Simpson index is relatively recent. Zhang & Zhou (2010) demonstrated that under independent sampling from a multinomial distribution,

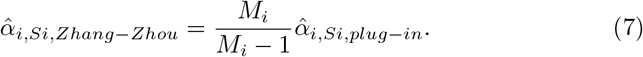

is unbiased and asymptotically normally distributed. However, since *M_i_* generally exceeds 1,000 in microbiome studies, the difference between the Zhang & Zhou (2010) and the plug-in estimate is negligible in our setting.

A number of approaches to estimating the Shannon index are also applicable to estimating the Simpson index. For example, Cao et al. (2017) investigate the performance of the zero-replace and low-rank approach to estimating the Simpson index. The extrapolation approach of Hsieh et al. (2016) also applies to the Simpson index. We compare our proposal, which we call `DivNet`, with these approaches in Sections 5 and 6.

#### 2.1.3. *α*-diversity with covariates

All of the estimates for *α_i_* discussed above are only functions of the abundance vectors **W***_i_*. Notably, none utilize the full abundance matrix *W* nor the covariate matrix *X*. Recently, Arbel et al. (2016) proposed a nonparametric Bayesian model that exploits structure in *W* as well as incorporating covariate information. However, the method is computationally expensive, and at present, an implementation only exists for *p* = 1. We compare our method to the method of Arbel et al. (2016) with respect to both estimation error and computation time in Section 5. We also note the recent method of Ren et al. (2017), which incorporates an error model into ordination methods, an alternative to diversity analysis in summarizing compositional data.

### 2.2. *β*-diversity

Similar to *α*-diversity, a large number of different *β*-diversity metrics exist, each highlighting different features of differences in ecosystems. Legendre & Legendre (2012, Table 7.2) provide a list of 26 *β*-diversity metrics along with some discussion. However, in comparison to *α*-diversity estimands, there exists almost no statistical literature on estimating *β*-diversity indices: estimating *β*-diversity indices is almost exclusively performed using plug-in estimators.

In general, small values of a *β*-diversity index indicate that the ecosystems have similar compositions, while large values indicate that the relative abundances differ between ecosystems, or that few taxa are shared by the ecosystems. This interpretation holds for both the Bray-Curtis and Euclidean indices discussed below.

#### 2.2.1 Bray-Curtis dissimilarity

The (observed) Bray-Curtis index (Bray & Curtis 1957) is defined as

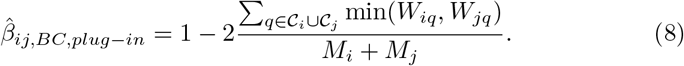

While we have not found any discussion of the target estimand in the literature, Eq. (8) suggests that

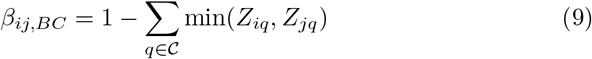

is the target estimand. Interestingly, in contrast to the other *β*-diversity indices discussed in the section, this estimate is not the MLE under a multinomial model.

While Arbel et al. (2016) focused on estimating *α*-diversity, because their method estimates the latent composition matrix *Z*, we also compare our proposed method to the estimate

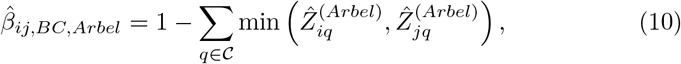

where 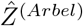 is the latent composition matrix estimate based on the procedure of Arbel et al. (2016).

#### 2.2.2 Euclidean distance

Finally, we mention the Euclidean distance between the relative abundance vectors,

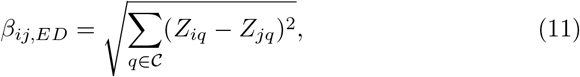

whose plug-in estimate is

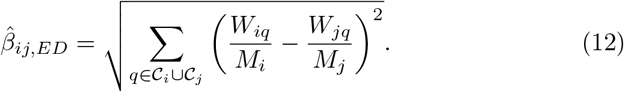

We are not aware of any other estimates for the Euclidean distance between relative abundances in the literature, but we will also compare to the estimate

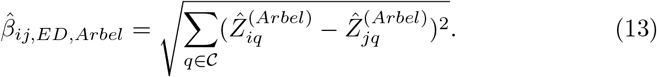

## 3. Estimating diversity in networked composition data

Members of ecological communities interact, displaying repeatable patterns in many different environmental settings (Faust & Raes 2012). For example, organisms may compete for resources, prey on each other, or cooperate in a symbiotic relationship. In the last decade, many methods have been developed to estimate the co-occurrence patterns of ecological communities, such as SparCC (Friedman & Alm 2012) and SPIEC-EASI (Kurtz et al. 2015). We will refer to co-occurrence patterns as *ecological networks*. As we show under simulation, ecological networks can have substantial effects on estimates of diversity. Here we propose an approach to estimating diversity in the presence of an ecological network. To our knowledge, this is the first method that explicitly accounts for co-occurrence patterns in diversity estimation.

### 3.1. Compositional data models

While the multinomial distribution is the canonical model for compositional data, the covariance between the number of observations in different categories is constrained to be negative. To deal with this issue, Aitchison (1982, 1986) developed the log-ratio model (see also Mandal et al. (2015)).This models the counts *W_iq_* as independent draws from a multinomial distribution,

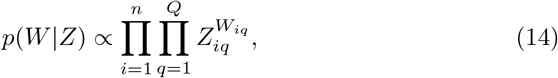

where *Z* ϵ *R^n^×^Q^* is a matrix-valued latent random variable that gives the underlying composition matrix for each of the samples: 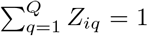 for all *i*. It then employs the log-ratio transformation by fixing a “baseline” taxon (taxon *D*) for comparison:

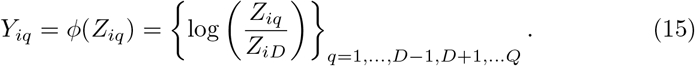

Note that the log-ratio transformation 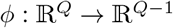 is invertible with inverse *ϕ*^−1:^.

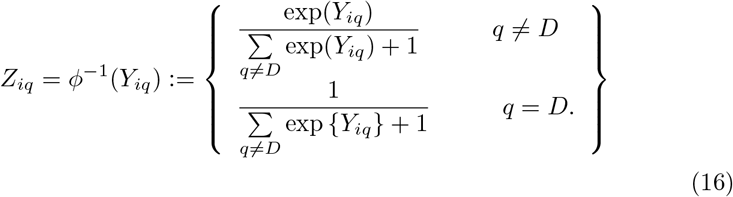

To permit exible co-occurrence structures between the taxa, the log-ratios are modeled by a multivariate normal distribution:

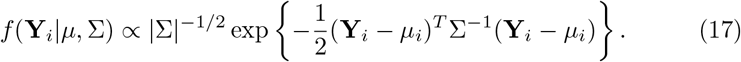

Finally, the mean of **Y***_i_* is linked to covariates via 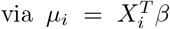, where 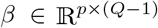. Under this model, *β_pq_* gives the expected increase in log 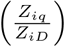 for a one-unit increase in *X_ip_*. For a discussion of the interpretation of this model on the scale of *Z_iq_*, we refer the reader to Billheimer et al. (2001).

### 3.2. Estimating diversity in the presence of a network

We propose using the log-ratio model to estimate *α*-diversity and *β*-diversity. Let 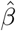 be an estimate of *β* under the log-ratio model. We discuss maximum likelihood estimators in detail in Section 4.1, and penalized maximum likelihood estimators in Section 4.2.

Suppose wish to estimate the *α*-diversity of an ecosystem with covariate vector 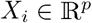. Define

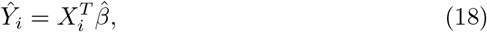

the expected value of the random variable **Y***_i_*, and define 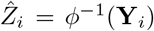, the fitted value of the latent composition. We then propose the following estimate of any *α*-diversity index 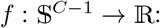

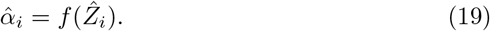

More explicitly,

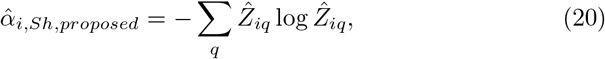

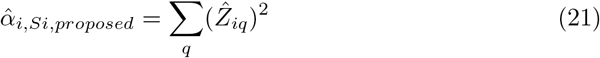

give our proposed estimates of the Shannon and Simpson indices. Similarly, for any *β*-diversity index 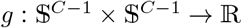, we propose

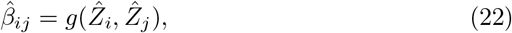

such as

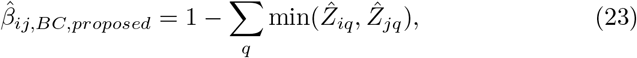

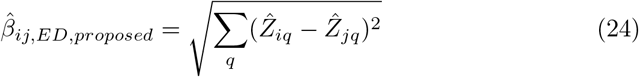

for the Bray-Curtis and Euclidean diversity indices. Note that if 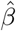 is the maximum likelihood estimate of *β*, then by invariance, the proposed estimates are the maximum likelihood estimates of the diversity indices.

This approach to diversity estimation has a number of key advantages not shared by other methods. Fundamentally, rather than describing a quantity associated with the sample (as is the case with plug-in estimates), the estimand is the diversity of the population from which the sample was drawn. This means that information is shared across all samples to obtain more precise and accurate estimates (see Section 5). In addition, samples *i* and *j* such that 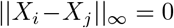 will have 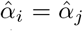 and 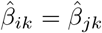 for any other sample *k*. In this way, biological replicates (samples where 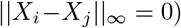 have equal diversity index estimates, in contrast to plug-in estimates and the estimates of Chao & Shen (2003), Hsieh et al. (2016), and Cao et al. (2017). Furthermore, we can use the model to estimate the diversity of ecosystems for which ecosystem survey data is not available but for which covariate information exists. While these advantages are shared with the method of Arbel et al. (2016), our method is substantially faster (Figure 3), and is available as an open-source R package with examples and tutorials illustrating its use.

## 4. Parameter estimation

### 4.1. Estimating model parameters

To estimate the parameter set *η* = (*β*, Σ), we consider a maximum likelihood approach. If *Y* were known, our optimization problem would be to find

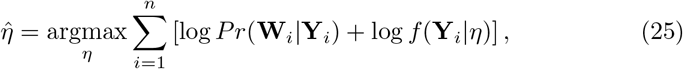

where

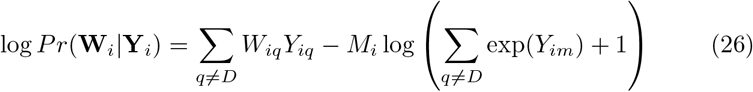

and

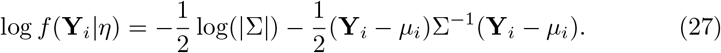

Alas, since *Y* is a latent random variable, we cannot directly optimize Eq. (25). Instead, we use the Expectation-Maximization algorithm (Dempster et al. 1977). The expected complete log-likelihood is

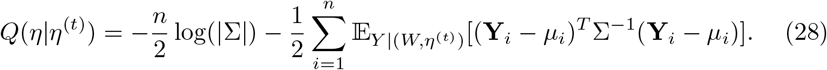

To estimate this expectation numerically, we follow Xia et al. (2013) and use the Metropolis-Hastings (MH) algorithm. Let 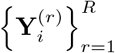 be *R* draws from the distribution of **Y***_i_*|**W***_i_*,*η*^(*t*)^. Given these draws, we can approximate theexpectation as follows:

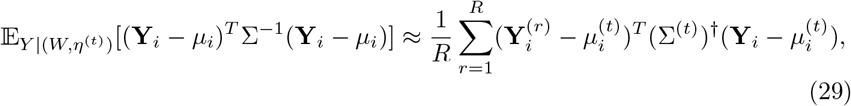

where † is the generalized inverse.

To generate the rth draw from *f* (**Y***_i_*|**W***_i_*, *η*^(*t*)^), we simulate a proposal 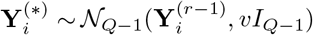, where *v* is a tuning parameter controlling the step size and *I_Q−1_* is the identity matrix of dimension *Q*−1. We then calculate the Metropolis acceptance ratio

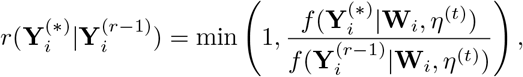

and simulate *u* ∼ Uniform(0,1). We set 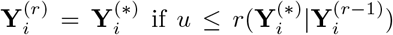, otherwise, we set 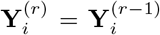. By initializing 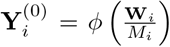, setting *v* = 0.01, and discarding the first 500 draws, we observe convergence to the target distribution on a variety of microbiome datasets, and acceptance ratios ranging 30-40%.

Having obtained an estimate of the expectation in Eq. (28), we turn our attention to maximizing *Q*(*η*|*η*^(*t*-1)^). Define *η*^(*t*)^ = argmax_*η*_ *Q*(*η*|*η*^(*t*-1)^). Given our draws 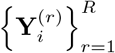 from *f*(**Y***_i_*|**W***_i_,η*^(*t*)^), our M-step of the EM algorithm gives the following estimates:

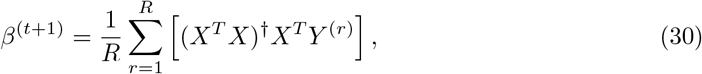

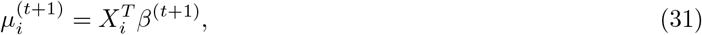

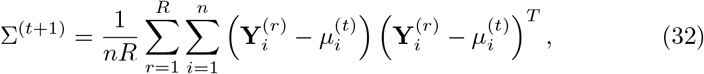

where 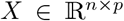 and 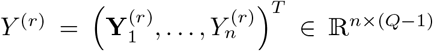. Inspection of convergence diagnostics (such as trace plots) on a variety of datasets indicates that *R* = 500 and 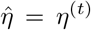 for *t* = 10 is generally sufficient to achieve stable estimates. We run the Metropolis-Hastings algorithm to approximate the distribution of **Y***_i_*|**W***_i_,η*^(*t*)^ in parallel over *i* = 1,…,*n* to reduce computation time. Our code is publicly available as an `R` package and can be found at github.com/adw96/DivNet.

### 4.2. Variance estimation

To test hypotheses about changes in diversity over environmental gradients it is necessary to have accurate estimates of the variance of the diversity estimates. These variance estimates can then be used in hypothesis testing (e.g., using the method of Willis et al. (2016)). We consider both parametric and nonparametric bootstrap approaches to estimating the variance of the diversity estimates produced by our model and evaluate them under simulation. For a given dataset (*W, X*), let 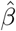 and 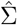 be the estimated values of *β* and Σ estimated by the algorithm described in Section 4.1.

The parametric bootstrap approach to estimating 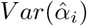 and 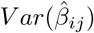 for arbitrary diversity indices works as follows: *B* datasets are simulated from the log-ratio model with 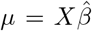 and 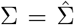. Then, for each of the *B* simulated datasets, bootstrap estimates 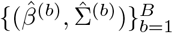 are obtained using the algorithm described in Section 4.1, and an estimate of the diversity index for sample *i* is obtained based on each simulated dataset (i.e., 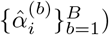). The parametric bootstrap estimate of 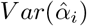 is then 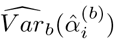, where 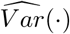 is the sample variance. An estimate of the variance of any *β*-diversity index can be obtained in the same way.

We also consider a nonparametric bootstrap approach to estimating the variance of our estimates. We uniformly at random select with replacement *n_sub_* elements from {1,…l,*n*} to obtain a set which we call 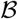. We then estimate 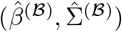 from 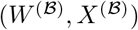, where 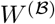 and 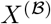 are the rows of *W* and *X* with row index in 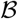, and use 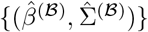 estimates to obtain 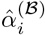. We repeat this process *B* times to obtain a set of estimates 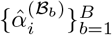 from which we calculate the non-parametric bootstrap estimate 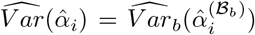 (and similarly for *β*-diversity).

The parameter Σ drives the variance in the log-ratio model: as 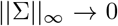, the distribution of *W* converges to a multinomial distribution. Therefore, the overdispersion of the log-ratio model relative to the multinomial model is driven by Σ. However, the number of taxa often greatly exceeds the number of samples obtained in microbiome surveys, and in this setting, (Σ^(*t*)^)^†^ may be a poor estimate of Σ^−1^ in Eq. (29), even for large *t*. We therefore consider replacing (Σ^(*t*)^)^†^ in Eq. (29) with a regularized estimate obtained from the graphical lasso (Friedman et al. 2008, Witten et al. 2011). Following the popular microbial network estimation software SPIEC-EASI (Kurtz et al. 2015), we use stability selection to select the regularization parameter (Liu et al. 2010, Kurtz et al. 2015). We also consider replacing (Σ^(*t*)^)^†^ with the maximum likelihood estimate restricted to the class of diagonal covariance matrices. Note that this approach to covariance estimation ignores variance attributable to inter-taxon interactions, but allows for overdispersion relative to the multinomial due to within-taxon interactions.

We evaluate the performance of these 6 approaches to estimating the variance of diversity indices (2 approaches to estimating the variance for each of 3 approaches to estimating the inverse covariance) under simulation. We design our simulation to mimic the dataset analyzed in Section 6, but with varying *Q*, the number of taxa and the size of the covariance matrix to be estimated. As is the case for the dataset of Section 6, we fix *p* = 1, *n* = 12, and set 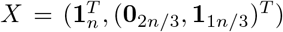. Let 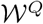 be the columns of the count matrix *W* of Section 6 corresponding to the *Q* most common taxa over all samples. Let 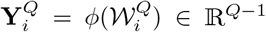, and 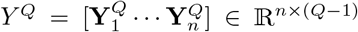. We set *β^Q^* = (*X^T^X*)^−1^*X^T^Y^Q^* and Σ*^Q^* to be the covariance of the columns of *Y^Q^ - Xβ^Q^*, and for each *Q*, we simulate data according to the log-ratio model with parameters *β^Q^*, Σ*^Q^* and *M_i_* = Σ*_q_ W_iq_*. Specifically, to simulate from the log-ratio model with parameters (*β*, Σ,*X,M*), we first simulate a matrix 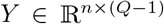 with *i*th row 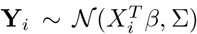, then calculate the matrix *Z* with ith row **Z***_i_* = *φ*^−1^(**Y***_i_*) (see Eq. (15)), and finally simulate the matrix 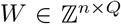 with **W***i* ~ Multinomial(*M_i_*, **Z***_i_*). Noting that *n* is small at *n* =12 (as is often the case for microbiome analyses), we choose *B* = 3 simulated datasets for the parametric bootstrap and *B* = 3 subsamples of size *n_sub_* = 6 for the nonparametric bootstrap approach.

We compare the estimated variance of the 6 methods in Figure 1 for a varying number of taxa *Q*. For brevity, only the variance of the Shannon index and Bray-Curtis index are shown. We observe that both parametric and nonparametric bootstrap variances are of similar magnitude, with parametric approaches generally having slightly lower median variance (left panels). In addition, to confirm that the estimated variance does not understate the true variance, we compare the difference between the estimated variance and the true variance for each method (right panels). The true variance of each method is estimated by repeatedly simulating data according to (*β^Q^*, Σ*^Q^, M*), estimating the diversity index for each simulated dataset and each covariance estimate, and calculating the variance of the estimated indices. We observe that the median difference between the true variance and the stated variance is near zero for the parametric approaches, but negative for the nonparametric approaches, indicating that nonparametric approaches tend to underestimate the true variance. However, none of the 3 approaches to covariance estimation show substantial advantage over the others. This suggests that the primary driver of variance in estimating diversity in microbial communities is within-taxon interactions (the diagonal elements of Σ), rather than between-taxon interactions (the off-diagonal elements of Σ). Given these results, we select the naïve (generalized inverse of the sample covariance) approach to estimating (Σ^(*t*)^)^−1^ as our default method. This approach is less computationally expensive than fitting the graphical lasso, while still permitting between-taxon interactions in the model. However, the functionality to estimate Σ via a structured approach is implemented in our `R` package.

**Fig 1.**
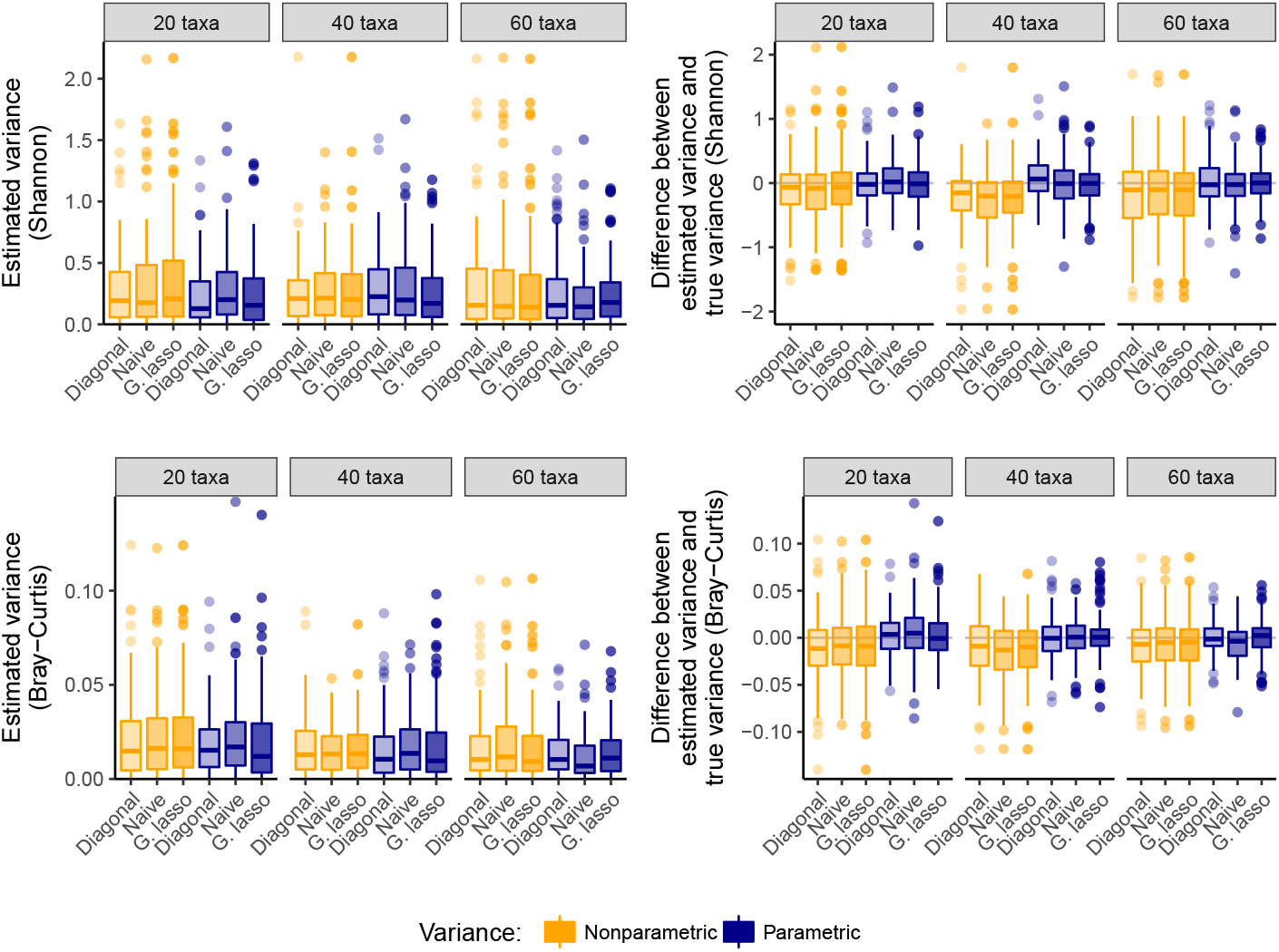
A comparison of candidate nonparametric and parametric bootstrap approaches to estimating the variance of diversity estimates under a model that incorporates microbial cooccurrence patterns. The parametric bootstrap has lower variance than the nonparametric bootstrap (left panel), and the median difference with true variance close to zero (right panel). No approach to covariance estimation consistently outperforms other approaches.

## 5. Simulation study

Having established estimators for diversity and variance, we now compare the performance of `DivNet` to estimates obtained from other methods. We simulate from the log-ratio model by specifying 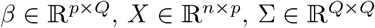, and 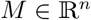 and simulating *W* as described in Section 4.2.

Note that the true relative abundance vector for sample *i* is 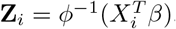, and so the true diversity indices can be calculated for each choice of *X* and *β*. For each of the 4 diversity indices that we consider in this paper (Shannon, Simpson, Bray-Curtis, and Euclidean), we obtain an estimate under the multinomial model and using the proposed estimation procedure. The procedure of Arbel et al. (2016) can be applied when *p* = 2, and so we set *p* = 2 and choose 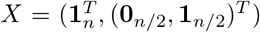 for all simulations. The `R` package `iNEXT` (Hsieh et al. 2016) applies to estimating Shannon and Simpson *α*-diversity indices, but not to estimating *β*-diversity indices. Note that many of the Shannon diversity estimates are almost identical to the Multinomial MLE for large values of *M* (*M* is commonly 10^5^ or greater in microbiome studies), including the estimates of Chao & Shen (2003) and Miller (1955), and for this reason we do not compare them here. For the same reason we also do not show the Simpson diversity estimate of Zhang & Zhou (2010). We use the `simulator` (Bien 2016) to manage the simulation study.

Throughout this section we evaluate *α*-diversity estimates using the mean square error (MSE) over all samples. The MSE of the kth simulation is

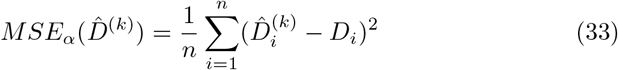

where *i* indexes the estimates for each of the *n* samples. We similarly evaluate the *β*-diversity estimates:

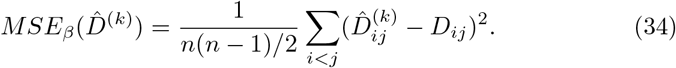

### 5.1. Estimation error decreases with sample size

We simulate the elements of 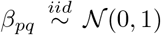. To construct Σ, we construct a matrix 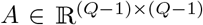 with elements drawn from a Uniform(−1,1) distribution, and construct a diagonal matrix *D* with diagonal elements forming an arithmetic sequence of length *Q* beginning at *σ_max_* and decreasing to a minimum of *σ_min_*. We then set Σ = *A^T^DA*. In this section we set *Q* = 20, *σ_min_* = 0.01, *σ_max_* = 5, and *M_i_* = 10^5^ for all *i*, and perform *K* = 200 simulations.

The performance of the proposed method for estimating diversity when data is simulated under this model is illustrated in Figure 2. For all values of *n* and all diversity estimands, the 25%, 50%, and 75% quantiles of 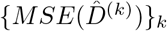 are uniformly lower for our proposed method compared to all other methods. The improvement is especially pronounced for the *β*-diversity indices.

**Fig 2.**
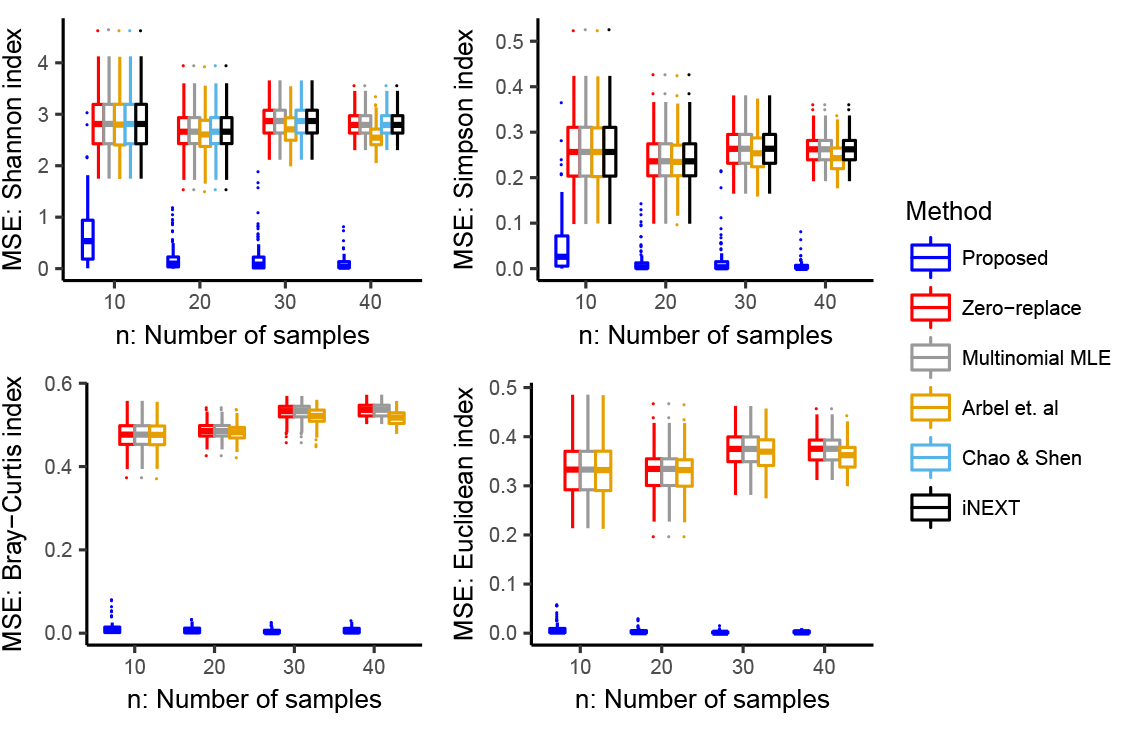
A comparison of the error of different estimators for *α*- and *β*-diversity for communities where the taxa are networked. When the network is ignored by the estimation procedure (e.g., Chao & Shen (2003), Hsieh et al. (2016) and the widely used “plug-in” estimate (multinomial MLE)), the error in estimating diversity can be substantial. The proposed estimation procedure, which specifically accounts for networks, outperforms other estimates for any sample size *n*. The distribution of mean squared errors (MSEs) is shown for 200 simulated datasets. In this simulation, there are *M* = 10^5^ microbes observed per sample, *p* = 2 predictors and *Q* = 20 taxa.

We find that the estimation error decreases as the sample size *n* increases for the proposed method and the method of Arbel et al. (2016), but not for
the Multinomial MLE and the iNEXT method (Figure 2). This is unsurprising, since neither the plug-in nor iNEXT estimates use information contained in the covariate matrix *X* in their estimates of diversity. Therefore, the additional information afforded by larger values of *n* is not leveraged by the plug-in nor iNEXT estimates, even when experimental replicates are available.

The results shown in Figure 2 are based on fitting our model with *t* = 10 EM steps and *r* = 200 MH draws per EM step. For these choices, we show computation time in Figure 3. Fitting our model with *t* = 10 EM steps and *r* = 200 is more computationally expensive than calculating the plug-in estimate, but less computationally expensive than fitting the model of Arbel et al. (2016) (with the default 10 chains) or using the package `iNEXT` (with default 40 knots and 50 bootstrap resamples). We note that our implementation leverages the `R` package `parallel` (R Core Team 2017) for parallelizing the MH algorithm employed at each E-step of the EM algorithm.

**Fig 3.**
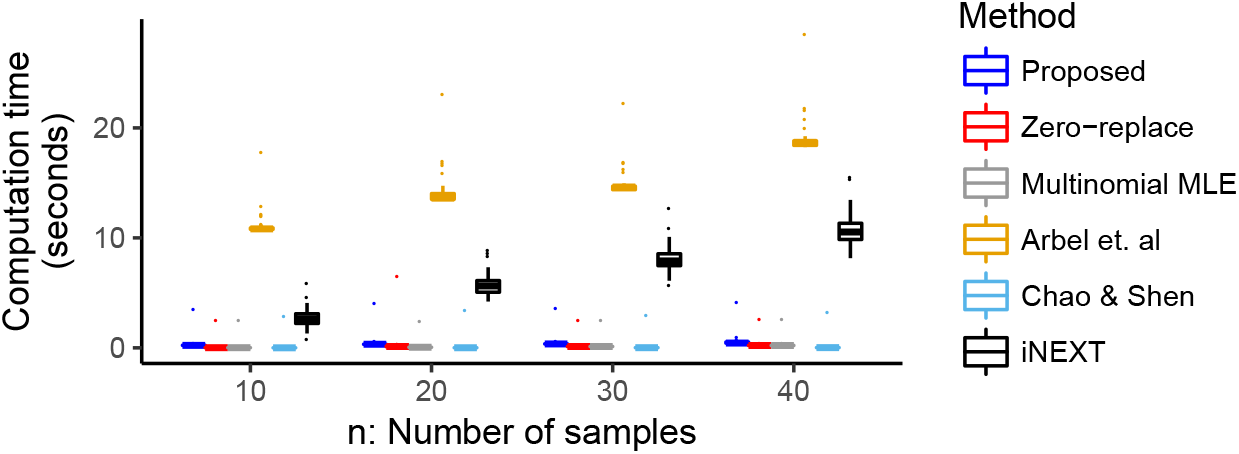
A comparison of the computing time of different estimators of diversity indices. Our parallelized EM-MH algorithm for estimation under a network model is competitive with closed-form estimates, and is substantially faster to compute than the rarefaction-extrapolation approach of iNEXT (Hsieh et al. 2016) and the nonparametric Bayesian approach of Arbel et al. (2016). The computation time of the 200 datasets used to produce Figure 2 is shown.

### 5.2. Estimation error is stable across networked communities

We now investigate the effect of varying the co-occurrence structure in the community on the estimation of diversity. Since larger values of the elements of Σ correspond to more strongly co-varying microbial abundances, by varying the elements of Σ, we can investigate the effect of the microbial concurrence network on diversity estimation. To vary the covariance structure in a systematic way, we vary σ*_max_*, the largest eigenvalue of Σ. We generate *β* and *X* as in Section 5.1, set *n* = 20, *Q* = 20, σ*_min_* = 0.01, *M_i_* = 10^5^ for all *i*, and perform *K* = 100 iterations for each choice of *σ_max_*. The results are shown in Figure 4. We see that estimating the diversity in microbial communities with strong occurrence structures is more challenging than estimating diversity in communities with co-occurrence structure similar to that of a multinomial model. However, the proposed method has lower MSE than all other methods that were investigated. Additionally, even when microbial abundances are simulated under a model with strong co-occurrence relationships, the proposed method can estimate the diversity with small MSE (Figure 4). In contrast, the estimation error increases as the co-occurrence relationships strengthen for all other methods. Co-occurrence relationships in microbial ecosystems are well documented (Faust & Raes 2012), indicating that a diversity estimation method tailored to networked ecosystems is of practical utility.

**Fig 4.**
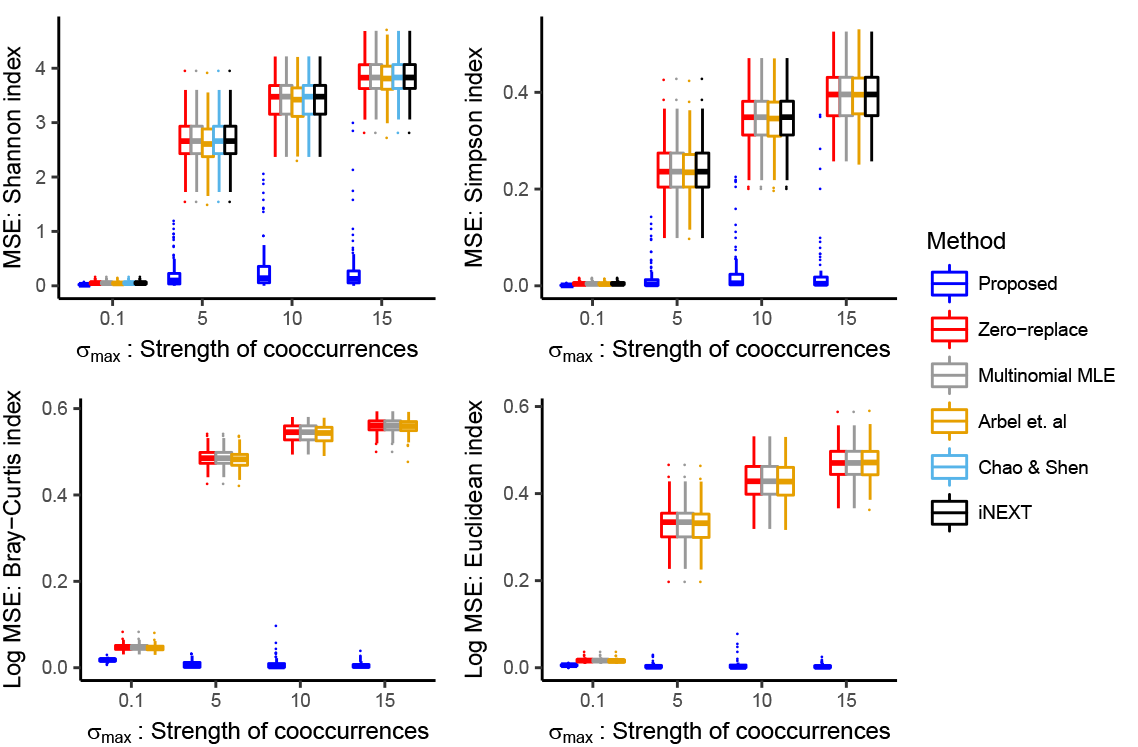
Diversity estimates that incorporate network structure dominate estimators that do not incorporate network structure in the presence of a strong co-occurrence network. However, network-based estimates perform well even when there is a very weak network structure. As *σ_max_* ⟶ 0, the network model converges to the multinomial model. However, we see that the proposed network model performs equally as well or better than estimates based on the multinomial model for all choices of ơmax. This appears to be the case for estimating both *α*- and *β*-diversity.

### 5.3. Estimation error is stable across large communities

Finally, since microbial communities often contain many taxa, we wish to confirm the performance of our estimator in large communities. In Figure 5, we see that the estimation error for the proposed method remains low even as the size of the community increases, while all other methods have increasing estimation error. In particular, we note that this is true even though the simulated communities are networked (*σ_max_* = 5), and the number of taxa exceeds the number of samples (*n* = 20). We therefore conclude that the procedure is appropriate for analyzing the diversity of communities with many taxa, such as microbial communities.

**Fig 5.**
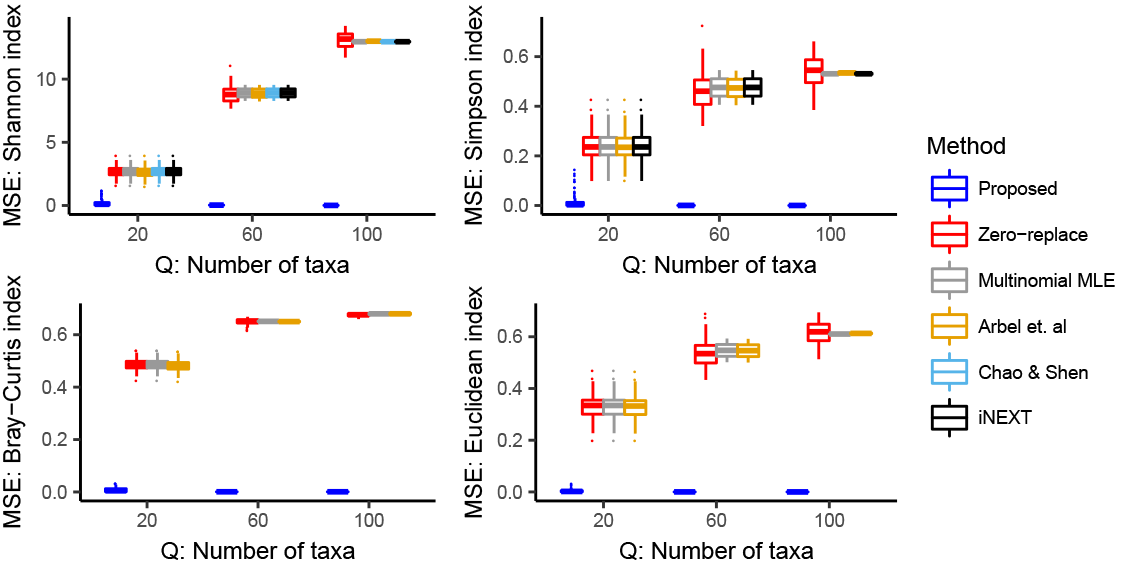
Diversity estimates that incorporate network structure dominate estimators that do not incorporate network structure over communities of any size. While most estimators have increasing error for larger communities, the proposed estimator’s error does not. In the simulation, we set *n* = 20 and *σ_max_* = 5.

## 6. Data analysis: Seafloor microbial diversity

Because of its coarse nature as a community-level summary, diversity analyses are especially relevant to studies of novel ecological communities. Lee et al. (2015) collected and analyzed microbial communities living on seafloor rocks on the Dorado Outcrop, an area of exposed basalt on the East Pacific Rise. Hydrothermal vents such as the Dorado Outcrop inform our understanding of microbe-mineral interactions in the subsurface. Samples were collected from the seafloor rock, including lithified carbonate (“carbonate,” *n* = 1), glassy, altered basalts (“glassy,” *n* = 4), and highly altered basalts (“altered,” *n* = 8). Analysis of the microbial communities on these rocks revealed 1490 distinct microbial taxa after filtering for low quality sequences (see Lee et al. (2015) and Lee (2018) for details surrounding sequencing and construction of the abundance table). Here we investigate if the community-level structure differs between the different rock types.

We investigate 30 choices for the *Q*-th taxon, whose abundance will be the denominator in the calculated log ratios. Since 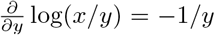 is smallest in absolute value for large *y*, we investigate the effect of setting *Q* to be a high abundance taxon. In particular, there were 66 amplicon sequence variants (ASVs) that were present in all samples, and so we uniformly at random select 10 ASVs from this collection of 66 ASVs, and compare the estimates of diversity obtained by setting each of these 10 taxa as the denominator taxon. We contrast these estimates with those obtained from ranging *Q* across the 10 most abundant taxa over all samples, i.e., let 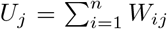, and call taxon *d* the *k*-th most abundant if *U*(*_k_*) = *U_d_* for *U*(*_k_*) the *k*-th order statistic of the {*U_j_*}*_j_*. We also compare 10 randomly selected taxa. The estimated Shannon, Simpson, Jaccard, and Euclidean diversities are shown in Figure 6 (right panels), indicating that, in practice, the diversity estimates are almost invariant to the choice of base taxon. Hereafter we select *Q* to be ASV 2 (a Nitrospirae of order Nitrospirales), which was the most abundant taxon that was observed in every sample.

**Fig 6.**
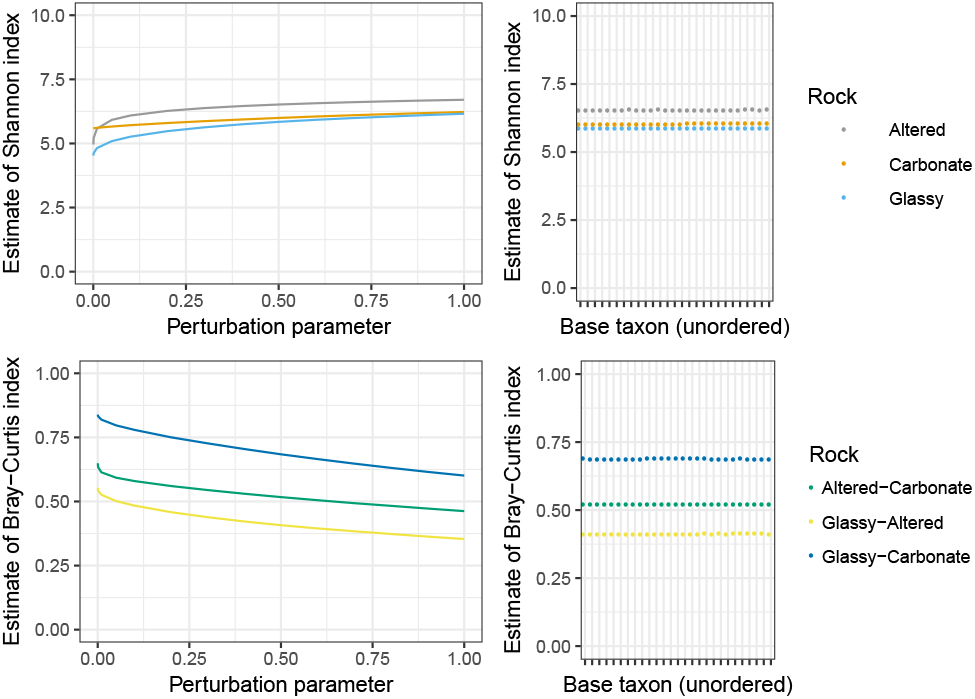
The log-ratio model described in Section 3 can only be fit to data with a minimum abundance greater than zero. Abundance data for microbiome studies is generally sparse, and 46% of the observed abundances of the Lee et al. (2015) dataset are zero. For this reason, it is common to add a perturbation offset *p* to the observed abundance table before fitting the log-ratio model. Here we see that the estimated diversity does depend on the choice of *p*.

In contrast to the stability of diversity estimates with varying *D*, we find that the effect of perturbing the zero counts can be substantial. As noted previously (Martín-Fernández et al. 2003, Cao et al. 2017), *W_ij_* is commonly zero for microbiome data, because many taxa do not occur in every sample (46% of the entries of our abundance table are zero). However *f* (*x, y*) = log(*x/y*) is only defined for *x,y* > 0, and so it is common to perturb the original abundance data *W* by adding a perturbation factor *p* ϵ (0,1) to create a new abundance table 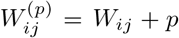, and the modeling the perturbed data *W*^(*p*)^. In Figure 6, we observe sizeable changes in the diversity estimates when varying *p* close to zero (at most 24%, −274%, −36% and −53% changes in Shannon, Simpson, Bray-Curtis, and Euclidean estimates for *p* = 0.001 compared to *p* = 0.5), but smaller changes when *p* is increased from 0.5 to 1 (at most 5%, −30%, −15% and −15% changes for *p* = 0.5 to *p* = 1). We therefore follow Cao et al. (2017) and choose *p* = 0.5 as the perturbation parameter for the remainder of our analysis.

Throughout this paper we have argued that the multinomial model is mis-specified for microbiome data. To investigate this claim for the dataset of Lee et al. (2015), we fit the log-ratio model and calculate the eigenvalues of of 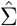. A test that that the largest eigenvalue of Σ is zero is rejected with *p* < 10^−5^ (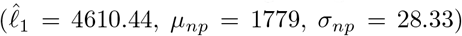 (Johnstone 2001, El Karoui 2003). This is strong evidence that a networked model is appropriate for this dataset.

Finally, we compare the estimates obtained from our method to the estimates obtained from other methods. Interval estimates are shown in Figure 7. While most methods produce similar estimates, we note a number of advantages of our proposal. Firstly, our method handles multiple covariates, which the method of Arbel et al. (2016) does not. Secondly, any diversity index that is a function of relative abundance can be estimated using our method, unlike the methods of Hsieh et al. (2016) and Chao & Shen (2003). Thirdly, our interval estimates are more symmetric around the median of the bootstrapped estimates compared to other estimates, indicating the greater stability of `DivNet` compared to other methods.

**Fig 7.**
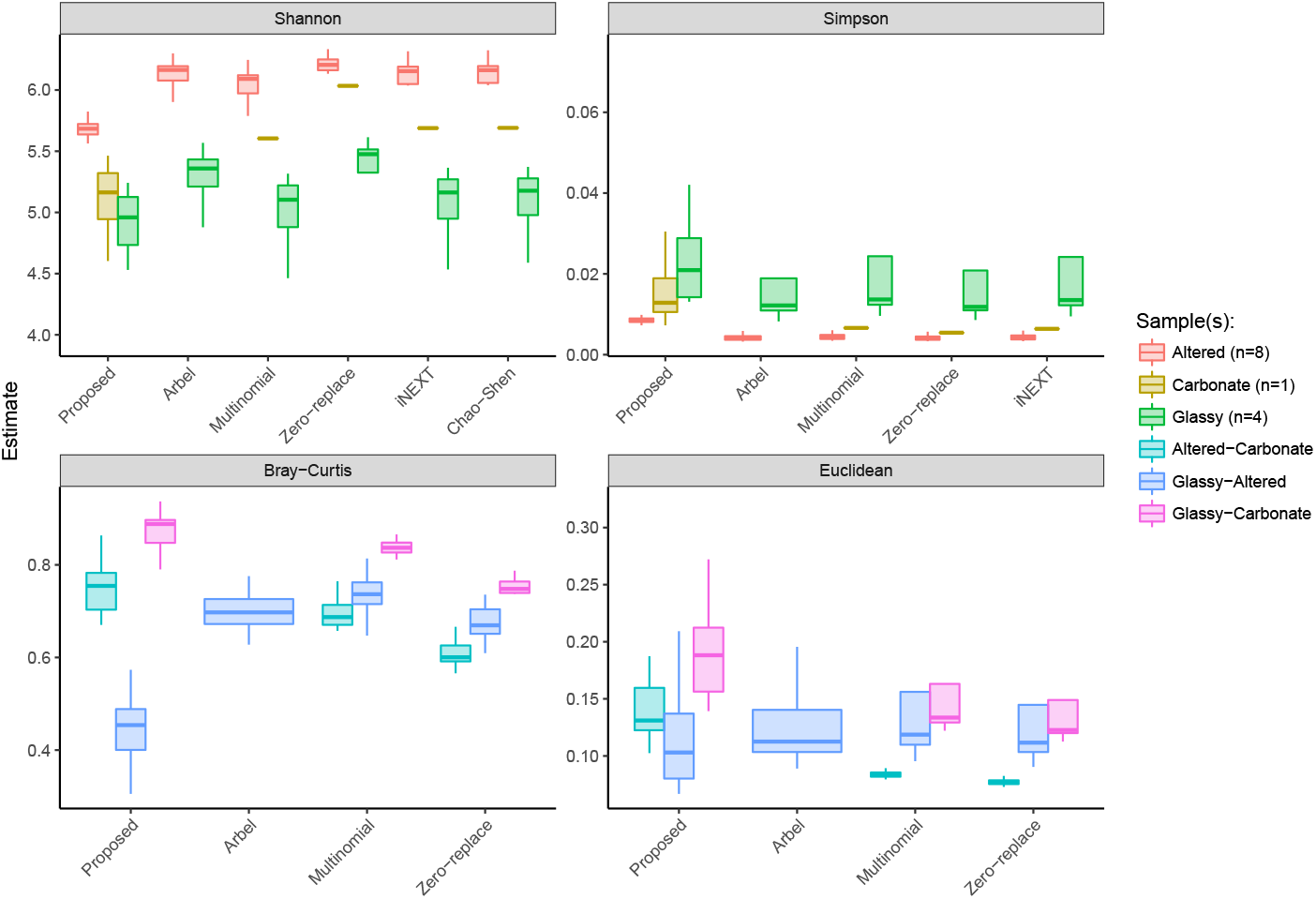
Lee et al. (2015) collected and analyzed microbial communities living on 3 types of seafloor basalts on the Dorado Outcrop. Here we compare a variety of estimators for 4 diversity indices (25% and 75% quantiles are shown). Our method works with multiple covariates, produces approximately symmetric interval estimates, and is the only method which can produce confidence intervals for the carbonate basalts.

The final advantage that we note is that for a sample condition which was only observed once (carbonate), an interval estimate is computable and of nonzero length. Our method is the only method that achieves this. The method of Arbel et al. (2016) cannot handle multiple covariates, and the remaining estimators can only produce point estimates based on a single sample. It is also worth noting that the interval for the sample observed once is wider than the interval for samples observed more than once (altered and glassy basalts), which is consistent with the amount of information available about this sample condition.

## 7. Discussion

Despite substantial evidence that strong co-occurrence networks exist in ecological communities, and a growing body of literature concerned with estimating cooccurrence networks, no methods that explicitly incorporate co-occurrence networks into diversity estimation currently exist. Here we propose a new method, called `DivNet`, to fill this gap. We have shown that `DivNet` is accurate, fast, performs well with a large number of taxa, and incorporates replicate and covariate information. It also permits extrapolation to experimental conditions that were not observed. It is available as a `R` package via github.com/adw96/DivNet.

By leveraging information from multiple samples, `DivNet` can estimate the relative abundance of a taxon in an ecosystem where it was not observed. However, a limitation of `DivNet` is it does not estimate the number of taxa that were missing in all samples. Therefore, when there are a large number of latent taxa, `DivNet` may miss the effects of these low abundance taxa. This weakness is shared by the estimators of Arbel et al. (2016) and Cao et al. (2016), while the estimators of Hsieh et al. (2016) and Chao & Shen (2003) adjust for missing taxa (but are only applicable to *α*-diversity). However, the latter 2 estimators cannot handle covariates nor repeated samples, which we believe significantly contributes to the strong performance of our method. We note that in the situation when no replicates or covariates are available, there are a large number of latent taxa, and *β*-diversity is not of interest, a practitioner may prefer these methods.

We suggest 3 avenues for further research that would build upon our proposed method. The first is to construct an estimator under the log-ratio model that estimates the number of missing taxa. However, this would require a principled approach to estimating the ecological network of a taxon that was not observed in any sample. A second avenue for research is to impose some structure, such as sparsity, on the relative abundance parameter *β*, whose dimension is large when there are a large number of taxa. Finally, since diversity indices that simultaneously incorporate relative abundance and phylogenetic information are commonly used by ecologists, extending the method to incorporate phylogeny is a challenging open problem.

All code to reproduce the simulations and data analysis, along with tutorials for using our package, are available at github.com/adw96/DivNet.

## Acknowledgements

The authors are grateful to Mike Lee for the dataset discussed in Section 6 and helpful discussions, to Daniela Witten for many insights on the model, and to Ali Shojaie and Erick Matsen for highlighting important references.

